# Inter-domain flexibility in F-Box proteins revealed by integrative modeling

**DOI:** 10.64898/2026.11.07.764542

**Authors:** Ben Shor, Fenglong Jiao, Huigang Shi, Haibin Mao, Clinton Yu, Merav Braitbard, Lan Huang, Ning Zheng, Dina Schneidman-Duhovny

## Abstract

F-box proteins, the substrate-recognition modules of the SCF (Skp1-Cul1-F-box) ubiquitin ligase complex, play a central role in targeted protein degradation by enabling the selective recruitment of diverse substrates. F-box proteins share a common architecture with a conserved anchor domain linked to the substrate-binding domain. Despite extensive structural characterization of F-box proteins, their potential conformational dynamics remain poorly understood. Here, we employ an integrative modeling approach combining cryo-EM structures, cross-linking mass spectrometry (XL-MS), and deep learning-based conformational ensemble prediction to profile the accessible conformations of F-box proteins. Our analyses reveal a significant inter-domain flexibility in FBXL2 and FBXL17 that enables sampling of a broad range of conformations within assembled SCF and CSN-SCF complexes. By contrast, other F-box proteins, including FBXL1 and FBXO22, exhibit intrinsic flexibility but are restricted to a considerably narrower conformational landscape. Together, our work highlights inter-domain plasticity as a tunable architectural feature of F-box proteins, which may shape their functional versatility within SCF complexes.

## Introduction

Cullin-RING Ligases (CRLs) represent the largest family of ubiquitin E3 ligases in eukaryotes, responsible for the targeted ubiquitination and subsequent proteasomal degradation of thousands of cellular proteins. The prototypical CRL1, or SCF (SKP1-CUL1-F-box) complex (1), achieves remarkable substrate diversity by utilizing interchangeable F-box proteins as specific recognition modules (2). These F-box proteins are structurally defined by an N-terminal F-box domain that anchors to the SKP1 adapter, and a variable C-terminal domain that recruits specific substrates (3). They are broadly categorized into three classes based on these C-terminal domains: FBXL (leucine-rich repeats), FBXW (WD40 repeats), and FBXO (other unclassified domains).

To maintain cellular homeostasis, the activity of the SCF complex is tightly regulated through a dynamic assembly cycle. Substrate recognition and complex assembly can occur sequentially: substrates first engage their cognate F-box protein, and the substrate-loaded receptor then assembles with the cullin scaffold. In the CUL1 system, this exchange of substrate receptors is mediated by CAND1, which acts as a substrate-receptor exchange factor (4–6). The COP9 Signalosome (CSN) serves as a gatekeeper in this cycle by catalyzing cullin deneddylation, since neddylated cullins cannot engage CAND1, whereas active, neddylated cullins drive substrate ubiquitination. Within this architecture, one side of the SCF complex must accommodate diverse substrates of varying size and shape, while the other side features the dynamic WHB domain of the cullin, which is regulated by CSN through deneddylation (7). The presence of such intrinsic flexibility on the cullin scaffold raises the question of whether comparable conformational dynamics exist on the substrate-recognition side, where the F-box protein must coordinate substrate capture under the steric constraints imposed by CSN regulation.

F-box proteins are commonly viewed as relatively rigid structural adaptors, supported by early X-ray crystallographic studies of isolated SKP1-FBX heterodimers. In contrast, an earlier study (8) used molecular dynamics simulations to suggest that CRL substrate-binding proteins possess intrinsic flexibility through an inter-domain hinge that orients the substrate for ubiquitin transfer. A thorough characterization of this flexibility, however, was limited by the availability of experimental structures and computational power at the time. Nevertheless, such a feature has been hinted at by select high-resolution cryo-EM reconstructions of ligase assemblies that exhibit less defined local density specifically within the F-box substrate-recognition regions (9). While complicated by sample heterogeneity or standard experimental variance during cryo-EM reconstruction, such localized resolution gradients can also be indicative of conformational dynamics.

In this study, we apply an integrative modeling approach (10, 11) to explore the potential inter-domain flexibility of the FBXL protein family. Through comparative structural analyses of newly resolved CSN-bound cryo-EM structural models alongside previously published ones, as well as generative deep learning-based conformational ensemble predictions (BioEmu (12)), we present evidence of conserved inter-domain flexibility across multiple F-box proteins. Using FBXL2 as a primary model, we further investigate these dynamics by employing comprehensive cross-linking mass spectrometry (XL-MS) coupled with integrative modeling (13). Together, these orthogonal approaches indicate that rather than acting as fixed platforms, FBXL proteins likely possess differential structural plasticity that allows them to sample a broader conformational landscape.

## Results

### Structural alignment reveals multiple conformations in FBXL proteins

Examining available FBXL structures, we found that FBXL17 adopts different conformations depending on its structural context. Aligning the F-box domain in the isolated SKP1-FBXL17 heterodimer bound to the KEAP1 BTB domain (PDB 6W66 (14)) with the fully assembled SCF^FBXL17^ complex bound to the BACH1 BTB domain (PDB 8UBT (15)), revealed that the substrate-binding domain occupies distinct conformations relative to SKP1 coupled with the N-terminal anchoring domain, with a Cα root mean square deviation (RMSD) of 17 Å (**Figure 1A**, orange and blue). A zoomed-in view of the inter-domain region in these two structures reveals that the displacement is concentrated at the short hinge connecting the F-box anchoring domain to the LRR substrate-binding domain, suggesting that this flexible linker is the source of the conformational change (**Figure 1B**).

**Figure 1.**
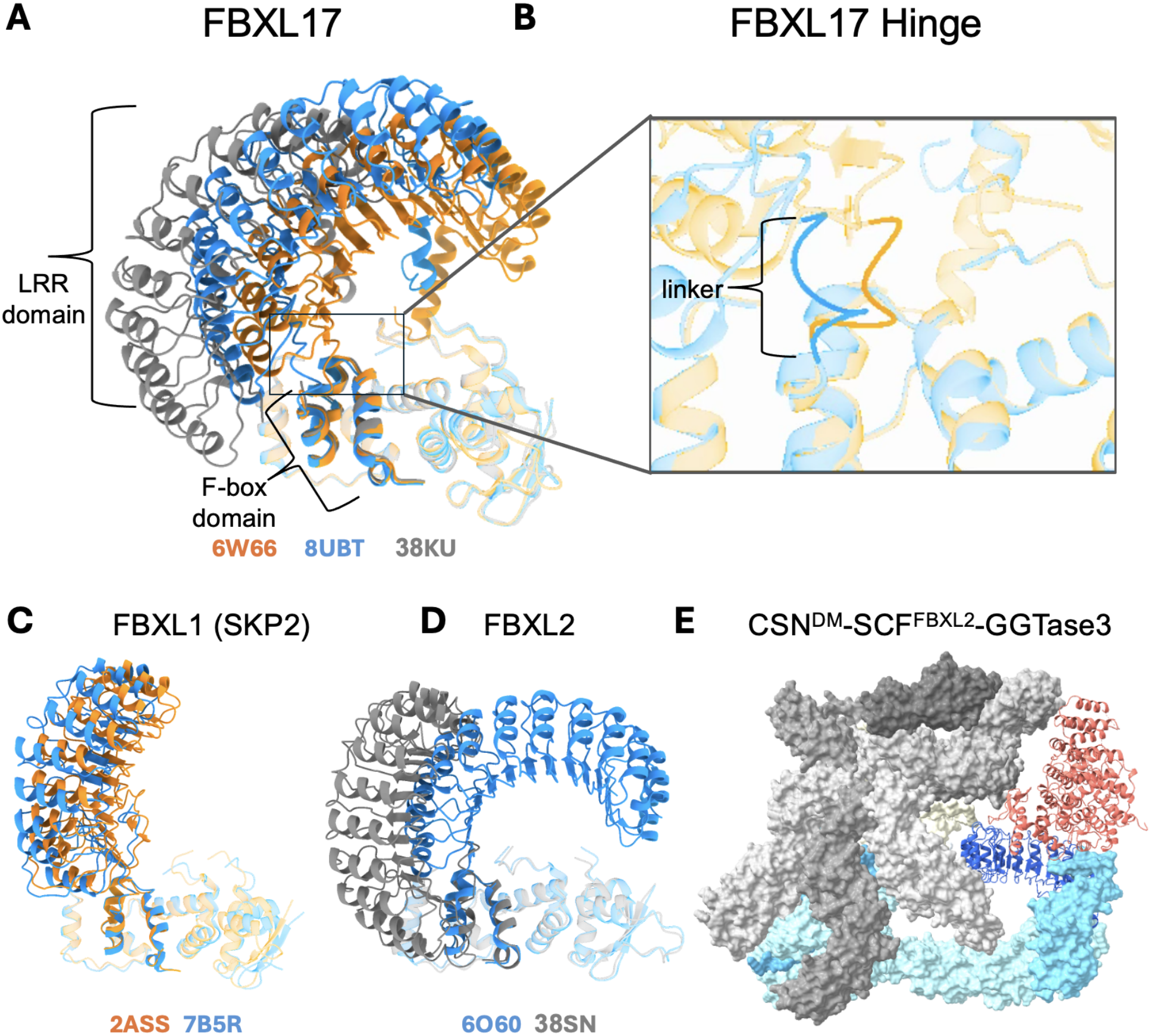
Structural alignments reveal inter-domain flexibility across the FBXL protein family members. Alignments in A, C, and D are anchored on the SKP1 subunit (shown as partially transparent) to highlight the relative movement of the FBXL substrate recognition domains (LRR domains). Structures were cropped from their original macromolecular contexts to display only the SKP1-FBXL modules. The source PDB entry of each structure is labeled in the matching color. **(A)** FBXL17: CSN^DM^-SCF^FBXL17^ complex aligned to the SCF^FBXL17^-BACH1 (PDB 8UBT) and SKP1-FBXL17-KEAP1 (PDB 6W66) complexes. **(B)** Zoomed view of the FBXL17 hinge in PDBs 8UBT and 6W66. **(C)** FBXL1: SCF^FBXL1^-Cks1 complex (PDB 2ASS) aligned to the Cul1-SCF^FBXL1^-Cks1-CDK2-CCNA2 complex (PDB 7B5R). **(D)** FBXL2: CSN^DM^-SCF^FBXL2^-GGTase3 complex (PDB 38SN) aligned to the SKP1-FBXL2-GGTase3 complex (PDB 6O60). **(E)** Cryo-EM structure of CSN^DM^-SCF^FBXL2^-GGTase3 (PDB 38SN). CSN^DM^ is colored in shades of gray, SCF in shades of blue, and GGTase3 in pink.

This observation motivated us to investigate whether the same phenomenon could be captured in a different physiologically relevant assembly. We therefore determined the cryo-EM structure of the inactive CSN (CSN^DM^) in complex with SCF^FBXL17^ at 4 Å resolution (**Supplementary Figure 1, Supplementary Table 1**). Strikingly, the substrate-binding domain of FBXL17 adopts a third, previously unobserved conformation in this context, with an RMSD of 21 Å and 33 Å to the previously published structures (**Figure 1A**). This reinforces the notion that FBXL17 remodels its architecture in a context-dependent manner.

To assess whether comparable inter-domain rearrangements are observed in other FBXL proteins, we searched the PDB (16) for additional family members with multiple deposited structures. Only one other FBXL protein, FBXL1 (SKP2), met this criterion, but with over 19 structures spanning more than five distinct biological contexts, providing a particularly well-sampled basis for evaluating conformational variability. Pairwise comparisons across these structures revealed that the substrate-binding domain adopts largely similar positions across contexts. The most pronounced difference was found between the SCF^FBXL1^-Cks1 complex (PDB 2ASS (17)) and the Cul1-SCF^FBXL1^-Cks1-CDK2-CCNA2 complex (PDB 7B5R (18)) (**Figure 1C**), where the substrate-binding domain RMSD, measured after alignment on the anchor domain, was only 4.5 Å, markedly smaller than the displacements observed for FBXL17 (up to 33 Å). Given the number and diversity of FBXL1 structures available, this comparison indicates that FBXL1 is less flexible than FBXL17, though some degree of conformational variability cannot be excluded. Together, these observations are consistent with a model in which the inter-domain hinge confers a degree of conformational freedom that is shared across FBXL proteins, but with considerably variable magnitude between family members.

To probe whether this flexibility extends beyond FBXL1 and FBXL17, we turned to FBXL2. Reasoning that its inter-domain flexibility might allow it to accommodate an additional binding partner within the CSN complex, we incorporated GGTase3, an FBXL2 modifying enzyme that is expected to occupy the substrate-binding site of the F-box protein, into the assembly and determined the cryo-EM structure of the resulting CSN^DM^-SCF^FBXL2^-GGTase3 complex (**Figure 1E, Supplementary Figure 2**). Aligning the FBXL2 subunit of this structure against the only existing SKP1-FBXL2 structure (PDB 6O60 (19)), which is likewise bound to GGTase3, revealed an even more pronounced inter-domain movement than in FBXL1 or FBXL17, with a substrate-binding domain RMSD of 43 Å between the two states (**Figure 1D**). While it is unknown whether CSN^DM^-SCF^FBXL2^-GGTase3 assembles under physiological conditions, the fact that such a complex can be stably formed and structurally resolved underscores the remarkable malleability of FBXL2.

Beyond these direct structural alignments, this intrinsic structural flexibility is further supported by the localized resolution profiles of our experimental EM maps. Since the models of CSN^DM^-SCF^FBXL2^ and CSN^DM^-SCF^FBXL17^ were built from averaged reconstructions, they correspond to structures in which particle-level conformational heterogeneity is averaged into a single state. Consistent with this, while the rigid CSN core resolved to approximately 4 Å, the local density corresponding to the SKP1-FBXL modules degraded to roughly 8 Å. This localized "fuzziness," which has also been noted in other structural studies of E3 ligase assemblies (9), is a known signature of conformational heterogeneity that cannot be fully captured by consensus refinement (20). It provides complementary physical evidence that the hinge region bridging the domains is highly flexible, and further motivates the orthogonal approaches (XL-MS and integrative modeling) used below to recover the underlying ensemble.

Overall, our analyses of the three distinct FBXL proteins (**Table 1**) suggest that conformational flexibility around the hinge connecting the anchor domain to the substrate-binding domain appears to be highly variable within the FBXL subfamily and such a feature may be an intrinsic and functionally relevant property of these SCF substrate receptors.

**Table 1.** FBXL structures that capture different states of the substrate recognition domain.

| FBXL member | PDB | Method / Resolution | Complex composition | Substrate binding domain RMSD |
|---|---|---|---|---|
| FBXL1 | 2ASS | X-ray (3.0 Å) | SCF<sup>FBXL1</sup>-Cks1 | 4.5 Å |
|  | 7B5R | Cryo-EM (3.8 Å) | Cul1-SCF<sup>FBXL1</sup>-Cks1-CDK2-CCNA2 |  |
| FBXL2 | 38SN | Cryo-EM (3.9 Å) | CSN<sup>DM</sup>-SCF<sup>FBXL2</sup>-GGTase3 | 43 Å |
|  | 6O60 | X-ray (2.5 Å) | SCF<sup>FBXL2</sup>-GGTase3 |  |
| FBXL17 | 38KU | Cryo-EM (3.7 Å) | CSN-SCF<sup>FBXL17</sup> | 17-33 Å |
|  | 8UBT | Cryo-EM (3.1 Å) | SCF<sup>FBXL17</sup>-BACH1 |  |
|  | 6W66 | X-ray (3.21 Å) | SCF<sup>FBXL17</sup>-KEAP1 |  |

Having focused on the FBXL family within CRL1, we asked whether comparable inter-domain motion might be a feature of substrate receptors in other CRL families. We performed the same alignment-based comparison on five receptors outside CRL1, spanning the CUL2, CUL3, CUL4, and CUL5 scaffolds, in each case aligning two deposited structures on the adaptor module analogous to SKP1 and measuring the displacement of the receptor domain (**Supplementary Figure 3**). Across these receptors, the receptor-domain RMSD ranged from roughly 11 to 17 Å. While this preliminary survey rests on a coarser analysis, it supports the possibility that adaptor-to-receptor flexibility is not confined to the F-box family but may be a recurring feature across CRL substrate-recognition modules.

### XL-MS datasets support the inter-domain flexibility of FBXL2

The static cryo-EM structures described above capture discrete conformational snapshots, but they cannot directly report on the dynamics of the FBXL proteins within an intact complex. To probe these dynamics in a more native context, we turned to cross-linking mass spectrometry (XL-MS), which is performed in solution and therefore captures distance constraints across a conformational ensemble under near-physiological conditions, rather than freezing a single state on a cryo-EM grid. Because FBXL2 displays the largest degree of structural flexibility in our prior analysis, we performed DSSO-based XL-MS analyses (21, 22) on four types of FBXL2-focused protein complexes with increasing compositional complexity: (i) the CSN holo-complex comprising the eight core subunits (CSN1-CSN8); (ii) the CSN-CRL complex containing Cul1, Rbx1, and Nedd8-conjugated Cul1; (iii) the CSN-CRL-SCF assembly additionally containing FBXL2 and SKP1Δ; and (iv) the complete GGTase3-bound complex further incorporating PTAR1Δ and RabGGTβ (**Supplementary Figure 4A, Supplementary Table 2**).

To validate the agreement between cross-links and solved structures, we started by focusing on highly reproducible cross-links identified in at least six independent experiments. Mapping these cross-links onto available three-dimensional structures of these complexes showed that the vast majority satisfied the expected distance constraint of 35 Å, supporting the validity of the XL-MS data (**Supplementary Figure 4B)**.

Next, we validated that the analysis can be expanded to include the interaction dynamics within these complexes. Our data analysis finds that four of the five identified CSN4-CSN6 cross-links were detected exclusively in apo CSN and were absent following CRL engagement. This was exemplified by the CSN6:K74-CSN4:K190 cross-link, whose Cα-Cα distance increased from 29.2 Å in apo CSN to 67.2 Å in the CRL-bound state. Similar increases in the corresponding Cα-Cα distances were measured for CSN6:K80-CSN4:K216, CSN6:K80-CSN4:K253, and CSN6:K74-CSN4:K216 residue pairs (**Supplementary Figure 4C)**. This observation is consistent with opening and reorganization of the CSN4-CSN6 interaction interface upon CRL binding described by previous studies (9, 23–25). To negatively control our observations, we examined the elongated Cul1 scaffold that is known to remain largely rigid throughout complex assembly. Indeed, most intra-Cul1 cross-links exhibited minimal distance changes across the different assembly states (**Supplementary Figure 4D**). We also acknowledge some limitations in our methods; for example, for CSN5, which goes through a conformational change between compositions, the mapped cross-link distances change, but the cross-links are satisfied in multiple conformations (**Supplementary Figure 4E**). Taken together, these results demonstrate the utility of comparative XL-MS for resolving both conserved and assembly-dependent interaction remodeling across these complexes, with emphasis on large inter-subunit plasticity.

To evaluate the structural dynamics of FBXL2, we mapped high-confidence cross-links (those observed in at least four of the six replicates) onto the CSN^DM^-SCF^FBXL2^ structure. The intra-protein cross-links had a satisfaction rate (< 35 Å) of 93% (129 of 138), but inter-protein cross-links performed significantly worse at 62% (60 of 96) satisfaction. The reduced satisfaction rate can be explained by the dynamics of inter-subunit interactions, which can not be captured in a single structure. Most notably, the satisfaction rate for cross-links between FBXL2 and the CSN core is 14% (only 1 of 7, **Figure 2A-B**). This sharp localized discrepancy indicates that a single, static cryo-EM structure does not adequately represent the true conformational ensemble of the FBXL2 subunit.

**Figure 2.**
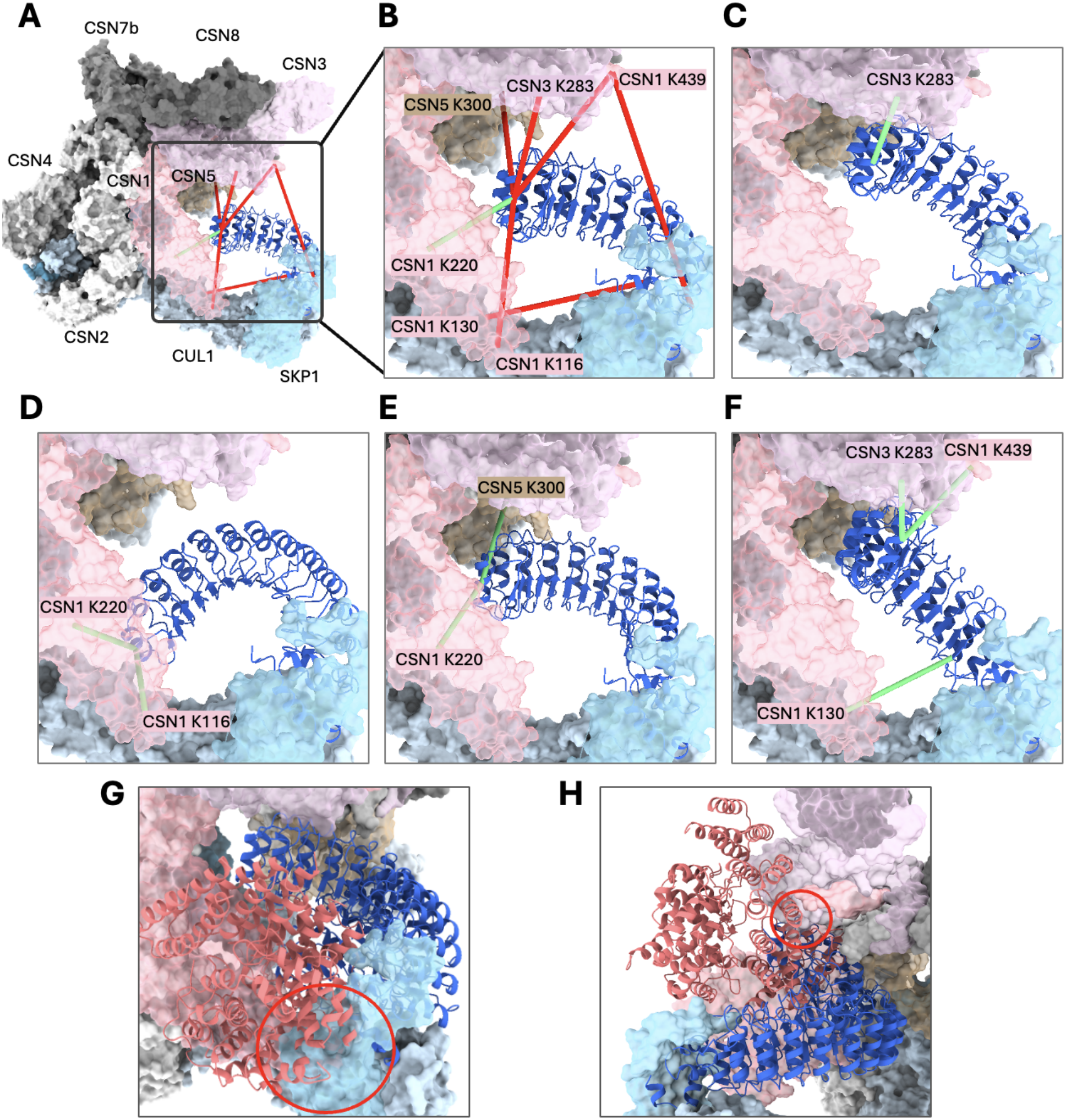
XL-MS-guided integrative modeling resolves violated cross-links by revealing a multi-state conformational ensemble for FBXL2. **(A)** The static, baseline cryo-EM model of the apo CSN-SCF^FBXL2^ complex. The CSN and SCF core subunits are displayed in surface representation, with the FBXL2 subunit shown in cartoon representation (royal blue). Thick red cylinders highlight high-confidence experimental cross-links between FBXL2 and the CSN core that are structurally violated (unsatisfied) by this single, rigid conformation, while the green cylinder indicates a satisfied cross-link. **(B)** Zoom view of FBXL2 and its cross-links in the static baseline structure. For each cross-link, its end in the CSN core is labeled with the residue identity. **(C-F)** Detail panels illustrating four distinct FBXL2 conformational states generated via PatchDock to account for the protein dynamics. Each discrete state successfully satisfies specific high-confidence inter-subunit cross-links (green lines) to various CSN subunits (CSN1, CSN3, CSN5) **(G-H)** PDB 6O60 superimposed onto FBXL2 conformational states generated via PatchDock, revealing steric clashes between the GGTase3 (salmon cartoons) and (G) SKP1 (light blue transparent surface) or (H) CSN1 (light pink transparent surface).

To computationally support this hypothesis, we employed integrative structural modeling to generate a dynamic, multi-state ensemble of FBXL2 within the CSN-SCF context. We performed protein-protein docking with PatchDock (26) to computationally re-dock the FBXL2 substrate-binding domain. The docking was driven by the experimental XL-MS distance restraints and the domain connectivity constraint to connect the FBXL2 anchor domain to the substrate-binding domain. This procedure yielded 8,248 candidate models, from which we selected those that best satisfied the cross-linking restraints while maintaining minimal deviation from the baseline cryo-EM structure (RMSD < 20 Å). The rationale behind this restriction is that we would like to prevent modeling large and low-probability conformational changes. Notably, no single sampled conformation could satisfy more than three of the seven cross-links without exceeding the 20 Å RMSD threshold.

We hypothesized that because no single conformation could satisfy a sufficient number of cross-links, capturing the protein’s true dynamics required a multi-state model where different conformations satisfy distinct cross-links. Testing this approach successfully generated a four-state ensemble, which increased the satisfaction rate of FBXL2-CSN cross-links to 86% (6 of 7 cross-links) (**Figure 2C-F**). This demonstrates that a single-state interpretation is biophysically insufficient; instead, FBXL2 likely needs to sample multiple structural states via its central hinge to satisfy the observed cross-linking network.

Having established this intrinsic flexibility in the apo state, we then asked what happens upon pseudo-substrate engagement. For this, we generated an XL-MS dataset for CSN-SCF^FBXL2^-GGTase3, which simulates an FBXL2 protein bound to a substrate and enables us to analyze how substrate engagement could potentially restrict this intrinsic flexibility. Indeed, evidence for reduced flexibility was raised as three of the seven high-confidence FBXL2-CSN cross-links observed in the apo state were completely absent in all six replicates of the GGTase3-bound composition.

We hypothesized that this cross-link absence is caused by steric occlusion, whereby the bulky substrate prevents FBXL2 from sampling its full conformational landscape without colliding with the surrounding complex. To test this, we aligned the previously resolved FBXL2-GGTase3 structure (PDB 6O60) with the specific states from our four-state apo model that account for these cross-links (**Figure 2D-E**). As predicted, superimposing the GGTase3 substrate onto these states resulted in severe steric clashes with the other subunits **(Figure 2G-H)**. This structural collision agrees with our conclusion that FBXL2 possesses intrinsic flexibility that is subsequently modulated and sterically restricted upon substrate binding.

Finally, we conducted a statistical analysis of crosslink consistency across experimental repetitions to serve as a biochemical proxy for localized structural rigidity. Our assumption is that crosslinks captured in a low number of experimental repetitions correlate with regions of high structural heterogeneity or transient, highly dynamic interactions. We define a relative index of flexibility as the ratio of inconsistent crosslinks (observed ≤ 2 times) to highly consistent crosslinks (observed ≥ 5 times) across each subunit of the CSN-SCF^FBXL2^ complex (**Figure 3**). When excluding subunits with fewer than 25 overall cross-links to prevent statistical noise, FBXL2 exhibits the highest ratio of inconsistent-to-consistent cross-links among all core components. This high degree of cross-linking variability provides robust, independent biochemical evidence that FBXL2 samples a broad and highly dynamic conformational space within the fully assembled complex.

**Figure 3.**
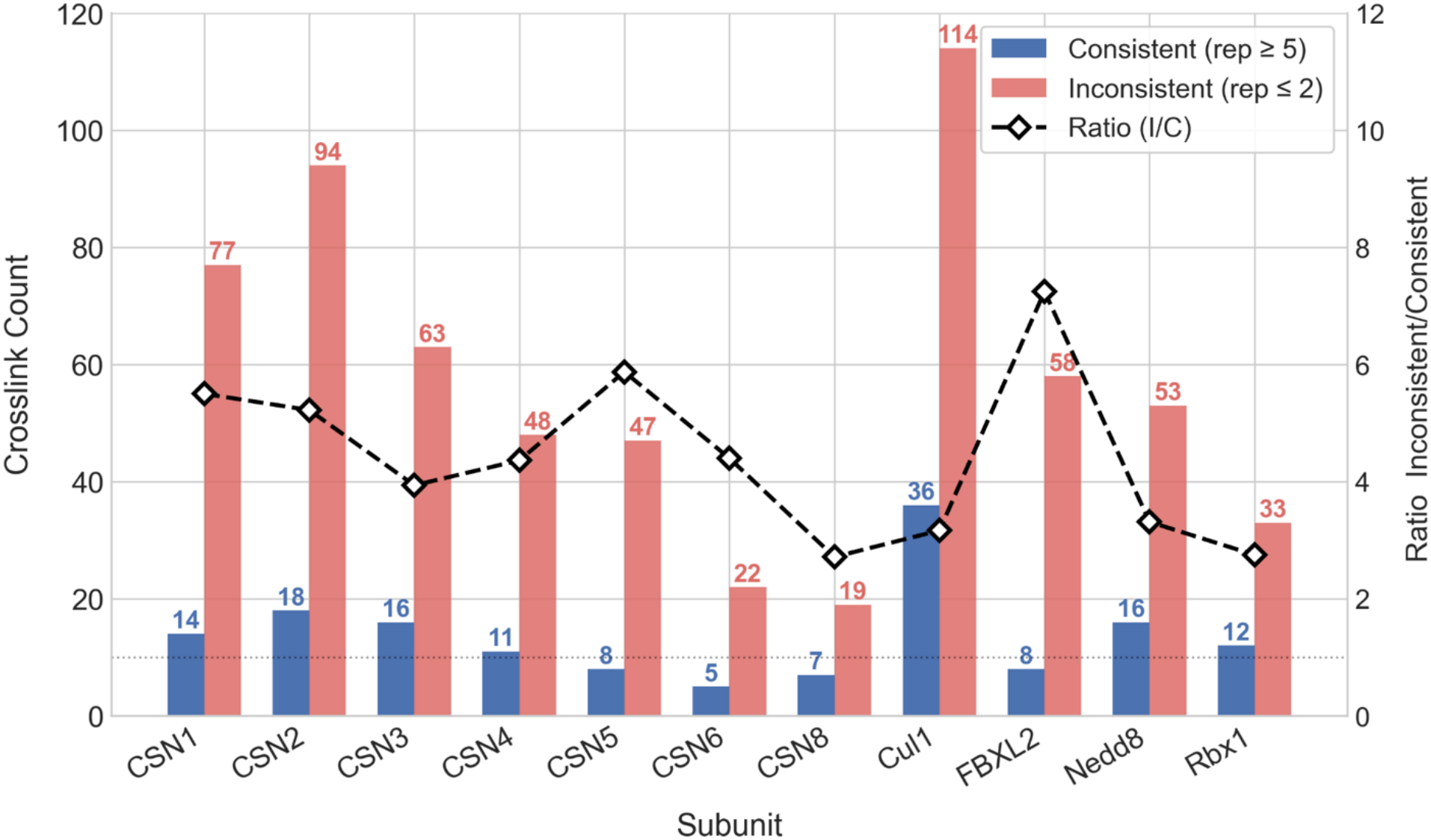
Cross-link consistency profiling highlights the intrinsic flexibility of FBXL2. Bar graph detailing the absolute counts of highly consistent cross-links (observed in ≥ 5 out of 6 replicates, blue bars) versus inconsistent cross-links (observed in ≤ 2 replicates, red bars) across individual subunits of the CSN-SCF^FBXL2^ assembly. The dashed black line represents the calculated ratio of inconsistent to consistent cross-links for each subunit, serving as a relative index of flexibility.

## Deep learning-based conformational sampling predicts flexibility across additional F-Box family proteins

Our experimental data establish FBXL2 as intrinsically flexible, and structural comparisons suggest that FBXL17 behaves similarly. To ask whether this dynamic behavior extends more broadly across the F-box family, we turned to computational ensemble prediction. We used BioEmu (12), an AlphaFold-based deep learning method that samples a distribution of plausible structural states from sequence input, enabling us to probe predicted conformational dynamics in F-box proteins for which multi-state experimental data are not available. For each complex, we sampled 100 structures and computed the median RMSD (mRMSD) that measures the median movement of the substrate-binding domain as a result of hinge flexibility (Methods).

We first asked whether BioEmu predictions agree with the extent of motion we observe experimentally for the three FBXL proteins with multiple available structures. For FBXL2, the predicted ensemble exhibited an mRMSD of 11.1 Å for the substrate-binding domain (**Figure 4A, gray**). The FBXL17 ensemble revealed an even larger mRMSD of 18.8 Å, with the sampled states consistent with all three of its experimentally resolved conformations, while also extending beyond them (**Figure 4B**). In contrast, BioEmu yielded a noticeably tighter ensemble for FBXL1, with an mRMSD of 7.6 Å (**Figure 4C**), substantially lower than the values obtained for FBXL2 and FBXL17, though not indicative of a fully rigid protein. BioEmu therefore correctly identified FBXL1 as the least flexible of the three, supporting its use as a comparative readout in cases where multi-state experimental evidence is not available.

**Figure 4.**
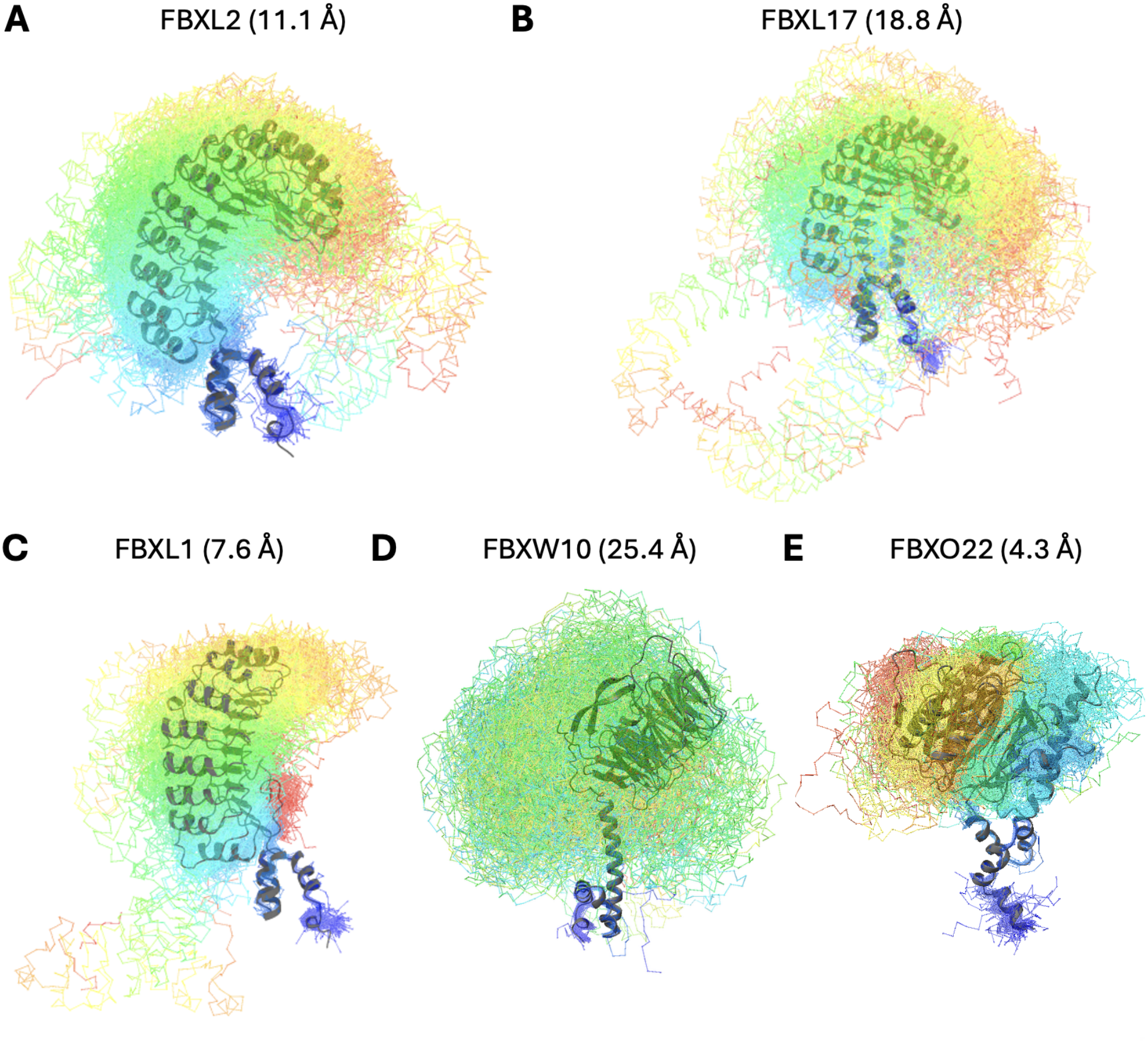
Deep learning-based conformational sampling of F-box proteins. For each complex, up to 100 structural states were sampled with BioEmu and superimposed on the anchor (SKP1-binding) domain to isolate the relative motion of the substrate-binding domain. Sampled conformations are shown as transparent rainbow chain traces, with the AlphaFold2 structure in gray (cartoons) for reference.

We next applied BioEmu to all F-box proteins (**Supplementary Table 3**). Within the FBXL subfamily, FBXL17 showed the highest mRMSD. Only two other proteins exceeded FBXL2, one of them being its paralog FBXL20, which had a comparable value (11.5 vs. 11.1 Å). Across the FBXW subfamily, nine members had relatively low mRMSDs between 4 and 9 Å, while FBXW10 stood out at 25.4 Å, which can be attributed to a long hinge that begins not at the SKP1-binding anchor but downstream of it (**Figure 4D**). The FBXO subfamily contained a notably higher fraction of highly flexible members: 10 of 34 proteins had mRMSDs above 10 Å. Closer inspection revealed, however, that in 9 of these 10, the substrate-binding-domain RMSD relative to its own averaged reference exceeded 4, a behavior seen in only 1 of the 30 FBXL and FBXW proteins. This suggests that, in this subset of FBXOs, the mRMSD is inflated by intra-domain conformational changes within the substrate-binding domain rather than by motion between the two domains. Although this mode of flexibility lies outside the scope of the present analysis, it provides additional evidence that conformational plasticity is a recurring feature of F-box proteins.

To experimentally challenge one of the BioEmu predictions, we focused on FBXO22, which (along with FBXW1) is one of only two F-box proteins outside FBXL1 and FBXL17 with more than five deposited structures, and which shows the narrowest experimental conformational spread of the two (up to ∼5 Å RMSD between FBXO22 structures, vs. up to 7 Å for FBXW1) and has an mRMSD of 4.3 Å (**Figure 4E**). To extend this comparison and test the prediction in the same experimental context used for our FBXL work, we determined the cryo-EM structure of CSN^DM^-SCF^FBXO22^ (**Supplementary Figure 5**). The new structure clustered with the prior FBXO22 structures (∼5 Å RMSD to the most divergent reference), indicating that the CSN-bound context, which revealed pronounced variability in FBXL2 and FBXL17, does not expose a substantially broader conformational range for FBXO22.

Taken together, the computational analysis refines, the conclusions from our experimental data: pronounced inter-domain hinge flexibility appears to be a relatively rare feature within the F-box family, with FBXL2 and FBXL17 standing out at the high end of the spectrum while most other members access only a more limited range of inter-domain conformations. This reframes hinge-mediated flexibility as a tunable architectural feature whose extent is modulated across F-box proteins, rather than a uniform, all-or-none property of the fold.

## Discussion

Here, we provide multi-modal evidence that certain F-box proteins, and in particular FBXL2 and FBXL17, are intrinsically flexible components of SCF complexes. Rather than behaving as rigid substrate-recognition scaffolds, these proteins appear to contain a mobile inter-domain hinge that connects the highly evolutionarily conserved SKP1/F-box anchoring module to the diverse substrate-binding domains. This conclusion is supported by structural comparisons, cryo-EM density, XL-MS datasets, and deep learning-based conformational predictions (**Figures 1-4**). Together, these approaches suggest that for these proteins, the substrate-binding domain can sample multiple positions relative to the SCF core.

Our experimental data support a model in which multiple FBXL2 states are required to explain the behavior of the complex. A single cryo-EM structure satisfies most cross-links across the CSN-SCF assembly, but fails specifically for FBXL2-CSN cross-links (**Figure 2B**). Cross-link-guided modeling resolves this discrepancy only when several FBXL2 conformations are allowed (**Figure 2C-F**). Given the sparse distance restraints, this four-state ensemble should be viewed as one representative realization rather than a unique structural solution. Importantly, the structure of the hinge does not suggest a small number of discrete, independently stable conformations. Rather, the simplest interpretation is that FBXL2 samples a continuous conformational landscape, with its intrinsic flexibility manifesting as apparent plasticity when different experimental approaches capture different states. This interpretation is further supported by computational conformational sampling, which predicts a similarly broad ensemble across many F-box proteins, without being provided any context.

Although our strongest experimental evidence comes from FBXL2, structural comparisons of FBXL17 reveal similar changes in the relative position of the substrate-recognition domain when aligned through SKP1, suggesting that pronounced inter-domain flexibility is shared between at least these two FBXL members. In these proteins, we suggest that inter-domain flexibility may represent a general architectural solution for SCF ligases: the conserved SKP1-CUL1-RBX1 core remains more stable, while the F-box substrate-binding domain retains enough freedom to accommodate substrates with different sizes, shapes, and degron geometries.

Our computational survey across the broader F-box family, however, suggests that the magnitude of this behavior varies considerably between proteins: the analyzed FBXW and FBXO members, as well as other FBXL proteins, generally show more modest predicted flexibility around the SKP1 anchor hinge, though none appear fully rigid. Consistent with this, our cryo-EM structure of CSN^DM^-SCF^FBXO22^ fell within the same narrow conformational range as previously resolved FBXO22 structures, without revealing a substantially different state. While these comparisons rely heavily on a single computational predictor and would benefit from further experimental validation, they suggest that intrinsic inter-domain flexibility may be a tunable rather than uniform architectural feature of F-box proteins, present across the family but exploited to differing extents by different members.

One reason such flexibility may have evolved is that SCF complexes face a fundamental geometric problem. The catalytic core is shared and relatively conserved, but substrates are highly diverse. A rigid substrate receptor would impose strong spatial constraints on which substrates could be recruited and how their lysines could be positioned for ubiquitination. A flexible F-box hinge could help to address this challenge by allowing the substrate-binding domain to explore multiple orientations while maintaining its connection to the SCF core. Such movement may help position different substrates for ubiquitin transfer, provide access to alternative substrate lysines, and allow the same ligase scaffold to engage structurally diverse targets. This may be one of several strategies that different F-box proteins use to accommodate their respective substrates, and proteins such as FBXL2 and FBXL17, which appear especially dynamic, may exploit this solution more than others. Of note, FBXL17 has been reported to have a large substrate repertoire in dimerization quality control (14, 27). FBXL2 is also known to ubiquitinate a variety of substrates, including both cytosolic and transmembrane proteins (28–32).

An additional contribution of this work is the way flexibility was inferred from features that are often treated as secondary observations. Poorly resolved or “fuzzy” cryo-EM density is commonly associated with local flexibility or conformational heterogeneity, but it is often discussed only as a limitation of the reconstruction rather than as mechanistic information. Similarly, XL-MS is widely used to provide distance restraints and to probe heterogeneous assemblies, but here we also used cross-link reproducibility across replicates as a subunit-level indicator of conformational variability (**Figure 3**). In this work, their combined use supports localized flexibility of the F-box protein within a complete CSN-SCF assembly.

These findings have important implications for current models of CSN substrate sensing. Rigid-body models often suggest that bulky substrates cannot be accommodated within CSN-bound SCF complexes, leading to the idea that CSN may sense substrate occupancy simply through steric exclusion. Our data suggest that this view is incomplete. If the F-box protein is flexible, then apparent clashes seen in static models may overestimate substrate incompatibility. Substrate-bound F-box proteins may still fit within CSN-SCF assemblies by adopting alternative conformations that are not captured by rigid docking.

At the same time, the substrate does not leave the system fully unrestricted. In the XL-MS dataset, GGTase3 binding removes several apo FBXL2-CSN cross-links, and structural modeling suggests that these lost contacts correspond to FBXL2 states that would clash with the bound substrate (**Figure 2G-H)**. Thus, substrate binding appears to limit the FBXL2 conformational landscape, but not necessarily to eliminate flexibility altogether. This leads to a revised model for CSN substrate sensing: CSN may not simply detect whether a substrate is present or absent. Instead, it may sense how substrate binding reshapes the conformational ensemble of the F-box protein.

## Methods

### Protein preparation and complex assembly

Human CSN was prepared by recombinant expression and reconstitution as described previously (25). Briefly, the CSN1-2-3 and CSN4-6-7 subcomplexes and CSN8 were expressed in *E. coli*, purified by affinity, ion-exchange, and size-exclusion chromatography, and assembled into the CSN seven-subunit complex. Recombinant CSN5^DM^ was expressed in ExpiSf9 insect cells (Thermo Fisher Scientific), purified separately, and incubated with the CSN7mer to reconstitute the complete CSN complex, which was further purified by ion-exchange and size-exclusion chromatography.

Human SCF complexes were prepared by recombinant expression and reconstitution as described previously (15, 19). Briefly, the FBXO22-SKP1, FBXL17-SKP1, and FBXL2-SKP1 complexes were co-expressed in Hi5 (*Trichoplusia ni*) insect cells. FBXO22-SKP1 was purified by glutathione-affinity chromatography, whereas FBXL17-SKP1 was purified by Ni-NTA affinity chromatography. Affinity tags were removed by TEV protease, followed by ion-exchange and/or size-exclusion chromatography. FBXL2-SKP1 and GGTase3, composed of PTAR1 and RabGGTB, were recombinantly expressed and purified by affinity chromatography followed by ion-exchange and size-exclusion chromatography as previously described.

## Cryo-EM sample preparation and data collection

For grid preparation, CSN^DM^ was mixed with the respective N8∼CRL1(neddylated CUL1), FBXL2-SKP1, FBXL17-SKP1, or FBXO22-SKP1 complexes at an approximately 1:1:1 molar ratio, with a final complex concentration of approximately 10 μM. For the CSN^DM^-N8∼CRL1-FBXL2-GGTase3 sample, the FBXL2-SKP1-GGTase3 complex was first assembled and purified, followed by incubation with purified CSN^DM^ and N8∼CRL1 to form the final complex. Fluorinated octyl maltoside (FOM) was added to all samples to a final concentration of 0.07% (w/v) immediately before grid preparation. Three microliters of each sample were applied to glow-discharged holey gold grids (Quantifoil R1.2/1.3, 300 mesh). The grids were blotted for 6 s at 10 °C and 100% relative humidity, plunge-frozen in liquid ethane using a Leica EM GP2 and stored in liquid nitrogen until data collection.

For the CSN^DM^-N8∼CRL1-FBXL2-GGTase3, CSN^DM^-N8∼CRL1-FBXL17, and CSN^DM^-N8∼CRL1-FBXO22 complexes, cryo-EM data were collected on Titan Krios transmission electron microscopes (Thermo Fisher Scientific) operated at 300 kV at the University of Washington and the HHMI Janelia Research Campus. Automated data acquisition was performed using SerialEM (33, 34) with a beam-image shift strategy and active beam-tilt compensation.

For the CSN^DM^-N8∼CRL1-FBXL2-GGTase3 dataset, images were recorded at a nominal magnification of 105,000× on a Gatan K2 direct electron detector operated in electron-counting mode, corresponding to a physical pixel size of 0.84 Å. Zero-loss energy-filtered images were acquired using a post-column Gatan BioQuantum GIF with a slit width of 20 eV. The dose rate was approximately 13.9 e^-^ per Å^2^ per second, with a total exposure of approximately 60 e^− Å^−2 fractionated into 60 frames.

For the CSN^DM^-N8∼CRL1-FBXL17 and CSN^DM^-N8∼CRL1-FBXO22 datasets, images were recorded at a nominal magnification of 105,000× on a Falcon 4 direct electron detector, corresponding to a physical pixel size of 0.743 Å. Zero-loss energy-filtered images were acquired using a Selectris energy filter with a slit width of 20 eV. Movies were recorded in electron-event representation (EER) format at a dose rate of approximately 10.1 e-per Å2 per second, with a total exposure of approximately 60 e-per Å2 per micrograph. Micrographs were collected over a nominal defocus range of -0.8∼3.0 μm.

## Cryo-EM data processing

All datasets were processed with cryoSPARC (35). The beam-induced motion correction and CTF estimation was conducted with cryoSPARC using patch motion correction and patch CTF estimation. The motion-corrected micrographs were curated by defocus, CTF fitting resolution (excluding >8 Å), and ice thickness. The curated micrographs were exported and subject to blob or template picking. After inspection of picked particles, the left particles were extracted with fourfold binning and subject to 2D classification. The selected particles were subjected to ab-initio reconstruction with 4 or 5 classes. All particles were then subjected to heterogeneous refinement with the reconstructions generated from ab initio reconstruction as reference. Particles of good reconstruction were used as templates to train a model for Topaz picking (36).

For CSN^DM^-N8∼CRL1-FBXL17, 150,929 good particles from 6,130 micrographs generated from sequential template picking, 2D classification were subjected to Topaz picking. 1,479,179 particles were picked, extracted and further classified via sequential 2D classification, ab-initio reconstruction and heterogenous refinement. 183,158 particles from good 3D classes were kept for non-uniform refinement (37) with C1 symmetry implemented. These particles yield a 3.0 Å reconstruction. To improve the density of the flexible FBXL17-SKP1 region, a binary mask was applied for non-alignment 3D classification. A total of 43,832 particles were selected and subjected to non-uniform refinement, yielding a reconstruction at 3.7 Å resolution.

The datasets for CSN^DM^-N8∼CRL1-FBXL2, CSN^DM^-N8∼CRL1-FBXL2-GGTase3, and CSN^DM^-N8∼CRL1-FBXO22 were processed using similar procedures. More detailed information can be found in Supplementary Figure 1, Supplementary Figure 2, and Supplementary Table 1.

## Model building and refinement

For the CSN^DM^-N8∼CRL1-FBXL2-GGTase3 and CSN^DM^-N8∼CRL1-FBXL2 structures, CSN^DM^-N8CRL1 (PDB 9EFM), CUL1-RBX1 (PDB 1LDJ), and FBXL2-SKP1-GGTase3 (PDB 6O60) were separately fitted into the 4.2 Å EM map using UCSF Chimera (38). The fitted models were further adjusted in Coot (39) using rigid-body refinement. The curated composite model was subsequently refined using Rosetta (40) and Phenix (41).

For the CSN^DM^-N8CRL1-FBXL17, and CSN^DM^-N8∼CRL1-FBXO22 structures, FBXL17-SKP1 (PDB 8UBT) and FBXO22-SKP1 (PDB 8UA6) were used as starting models where applicable. The composite models were manually adjusted in Coot and sequentially refined using Rosetta and Phenix. Structural figures were prepared using PyMOL(42) and UCSF ChimeraX(43).

## Cross-linking of CSN complexes and SEC fractionation

The CSN-CRL complex was assembled by incubating Nedd8-Cul1-Rbx1 with the CSN mutant at a 1:1 molar ratio on ice for 30 min prior to cross-linking. The CSN-SCF and CSN-SCF-GGTase3 complexes were assembled by first mixing Nedd8-Cul1-Rbx1 with either SKP1-FBXL2 or SKP1-FBXL2-GGTase3 on ice for 15 min, followed by incubation with the CSN mutant at a 1:1 molar ratio on ice for 30 min before cross-linking. DSSO cross-linking was performed on CSN, CSN-CRL, CSN-SCF, and CSN-SCF-GGTase3 complexes at a protein-to-linker ratio of 1:250 for 1 h at room temperature, with three biological replicates (22). To further optimize cross-linking and reduce conformational dynamics, we tested additional conditions, including a lower linker concentration (1:100), reduced temperature (on ice), and mild pre-fixation with low concentrations of formaldehyde (FA; 10:1 lysine:FA ratio). In total, four cross-linking conditions were evaluated (1:250 RT, 1:100 RT, 1:100 on ice, and 1:100 with FA pre-treatment), yielding six replicates overall. After cross-linking and quenching with 50 mM NH4HCO3 for 15 min, cross-linked complexes were transferred onto a 30 KDa FASP centrifugal filter and washed with 25mM NH4HCO3. The proteins were reduced by 3.3 mM TCEP for 30 min and alkylated with 16.5 mM iodoacetamide for 30 min in dark. After washing with 25 mM ammonium bicarbonate, proteins were reconstituted in 60 µL of 8 M urea in 25 mM ammonium bicarbonate. Lys-C (enzyme to protein ratio of 1:100) was added to the solution and the mixture was incubated at 37 °C for 4 h. Then, the concentration of urea was reduced to 1.5 M for trypsin digestion (enzyme-to-protein ratio of 1:50) at 37°C overnight. The peptide digests were extracted, desalted by Sep-Pak C18 cartridge and stored in -80°C before SEC enrichment. Cross-linked peptides were dissolved in 30% ACN/0.1% TFA and fractionated on a Superdex™ 30 Increase 3.2/300 column with an Agilent 1260 Series HPLC. The three fractions with inter-linked peptides (F22-24min, F24-26min and F26-28min) were collected and dried for LC-MSn analysis(22).

## LC-MSn analysis

Cross-linked peptides were analyzed by LC-MSn using an UltiMate 3000 RSLC coupled with an Orbitrap Fusion Lumos mass spectrometer. Samples were loaded onto a 50 cm x 75 μm Acclaim PepMap C18 column and separated over a 180 min gradient of 4% to 25% acetonitrile at a flow rate of 300 nL/min. Ions with charge of 4+ to 8+ in the MS1 scan were selected for MS2 analysis. The top 4 most abundant fragment ions in MS2 scan were further fragmented by CID. MS1 and MS2 scans were acquired in the Orbitrap whereas MS3 scans were detected in the ion trap. For MS2 scans, the resolution was set to 30,000, the AGC target 1e5, the precursor isolation width was 1.6 m/z, and the maximum injection time was 70 ms for CID. The CID-MS2 normalized collision energy was 23%. For MS3 scans, CID was used with a collision energy of 35%, the AGC target was set to 2e4, and the maximum injection time was set to 150 ms.

## Identification of Cross-linked Peptides

MS3 spectra was extracted by PAVA (UCSF) and subjected to Protein Prospector (v.6.3.5) (44) for database searching (using Batch-Tag against the database containing the 15 proteins). The mass tolerances were set as ±20 ppm for parent ions and 0.6 Da for fragment ions. Trypsin was set as the enzyme with three maximum missed cleavages allowed. Cysteine carbamidomethylation was set as static modifications. A maximum of four variable modifications were also allowed, including methionine oxidation, N-terminal acetylation, and N-terminal conversion of glutamine to pyroglutamic acid. Three defined DSSO cross-linked modification on uncleaved lysines, including alkene (C3H2O, +54 Da), thiol (C3H2SO, +86 Da) and sulfenic acid (C3H4O2S, +104Da) were also selected as variable modifications. The in-house software XL- Tools was used to automatically identify, summarize and validate cross-linked peptides based on Protein Prospector database search results and MSn data.

## Integrative Structural Modeling with PatchDock

To generate a dynamic, multi-state conformational ensemble of FBXL2 within the apo CSN-SCF complex, integrative protein-protein docking was performed using the PatchDock online web server (26). The rigidly defined CSN core and the SKP1-anchor region of the newly resolved cryo- EM model were held constant, while the C-terminal substrate-binding domain of FBXL2 was decoupled and computationally redocked. Experimental XL-MS distance measurements were utilized to guide the docking and subsequent filtering. The initial PatchDock run generated 8,248 distinct docked configurations. To assemble a biologically plausible multi-state model, these configurations were systematically filtered based on a dual-criteria approach: (1) the ability to satisfy high-confidence inter-subunit cross-links (observed in ≥ 4 replicates) between FBXL2 and the CSN that were violated by the static cryo-EM model, and (2) structural proximity to the baseline experimental state, enforced by a root-mean-square deviation (RMSD) threshold of < 20 Å. A final four-state ensemble was selected that maximized the cross-link satisfaction rate to 86%. For the subsequent steric hindrance analysis, the experimental coordinates of the FBXL2- GGTase3 complex (19) (PDB 6O60) were structurally aligned to the individual states of this generated apo ensemble to evaluate potential volumetric clashes between the GGTase3 substrate and the surrounding CSN-SCF architecture.

## Conformational Ensemble Prediction

Reviewed human entries of the FBXL, FBXO, and FBXW subfamilies were retrieved from UniProt (45) (gene-name wildcards FBXL*, FBXO*, FBXW*; Homo sapiens; reviewed) and filtered to those carrying an annotated F-box domain feature. For each entry, we retrieved the canonical sequence, the F-box residue range, any annotated substrate-domain repeats (LRR, WD40, etc.), and the corresponding AlphaFold2 model from the AlphaFold Database (46). For FBXO proteins, which have substrate-binding domains that are harder to characterize and compare, we only kept proteins with an experimentally resolved structure in the PDB.

Each AF2 model was trimmed to retain the F-box and the C-terminal substrate-binding fold. The N-terminus was anchored to the start of the F-box annotation, with up to about five residues of upstream extension allowed when they were ordered, and with explicit clipping of any folded N- terminal accessory domains (for example, the hemerythrin domain in FBXL5, or the JmjC, CXXC, and PHD modules in the KDM2A/B-bearing FBXLs). The C-terminus was placed at the end of the most C-terminal run of at least 20 consecutive residues with pLDDT greater than 70, then extended outward to cover annotated substrate-domain repeats when present, and a bridge-and- extend rule that captured short ordered tails: contiguous clusters of at least 5 residues with pLDDT of at least 80 within 30 residues of the boundary, bridging any short low-pLDDT gap. Low-pLDDT residues inside the final trim range were preserved for sequence continuity but flagged in the per- protein metadata so downstream metrics and alignments could exclude them.

For each trimmed protein, conformational samples were generated with BioEmu (target 100 conformers; 75 to 97 obtained per FBXL protein). Each sample was rigid-body aligned to a reference conformation using BioPython Superimposer on the Cα atoms of the F-box "anchor core", defined as positions 10 through 40 within the F-box annotation, excluding any low-pLDDT residues. The first ten F-box residues were omitted from the alignment set because they are partially disordered in many family. The resulting transform was then applied to every atom in the sample.

A protein’s flexibility was scored as the mean RMSD to the ensemble mean of the substrate- binding-domain Cα atoms, where the substrate-binding domain was defined uniformly as all trimmed residues not in the F-box, excluding low-pLDDT residues. For each protein, every BioEmu sample was first brought into a common coordinate frame by superimposing its F-box anchor Cα atoms (positions 10 to 40 within the F-box annotation) onto those of the first sample. The per-atom average of the substrate-binding-domain Cα coordinates was then computed across the ensemble and used as the reference, and for each sample the substrate-binding- domain Cα RMSD to that reference was recorded. The reported flexibility is the median of these per-sample RMSDs across the N samples.

## Data availability

MS raw data have been deposited in PRIDE (Project accession: PXD081048; Password: z7WBKCOybEvh). The coordinates and cryo-EM maps were deposited in the Protein Data Bank (PDB) and the Electron Microscopy Data Bank (EMDB) with the following accession numbers: CSN^DM^-N8∼CRL1-FBXL2: pdb_000038qo (PDB 38QO), EMD-79052; CSN^DM^-N8∼CRL1-FBXL2- GGTase3: pdb_000038sn (PDB 38SN), EMD-79111; CSN^DM^-N8∼CRL1-FBXL17: pdb_000038ku (PDB 38KU), EMD-78892; CSN^DM^-N8∼CRL1-FBXO22: pdb_000038xz (PDB 38XZ), EMD-79223.

## Acknowledgements

This work was supported by National Institutes of Health grants R35 GM145249 to L.H, ISF grant 737/23 and ERC grant 101171306 to D.S. N.Z. is a Howard Hughes Medical Institute Investigator.

B.S. is supported by the Clore PhD fellowship. The authors would like to thank R. Yan, Z. Zhang, N. Spellmon, and Z. Yu at the Cryo-EM Facility on the Janelia Research Campus of the Howard Hughes Medical Institute, and W. Jiang and J.D. Quispe at the Arnold and Mabel Beckman Cryo-EM Center at the University of Washington for their assistance in electron microscopy data acquisition.

## Author Contributions

D.S., L.H., and N.Z. conceived the project. H.M. and H.S. constructed and purified all protein samples in this study. H.S. performed all cryo-EM grid preparation, specimen screening, data collection, and processing tasks. F.J. performed cross-linking mass spectrometry experiments and data analysis. C.Y. assisted on mass spec data acquisition. B.S., M.B. and D.S. analysed the in vitro cross-linking mass spectrometry data. B.S. performed ensemble prediction experiments and multi-state modeling. The manuscript was written by B.S., F.J., H.S., and D.S., with input from all authors.

---

**Supplementary Figure 1.**
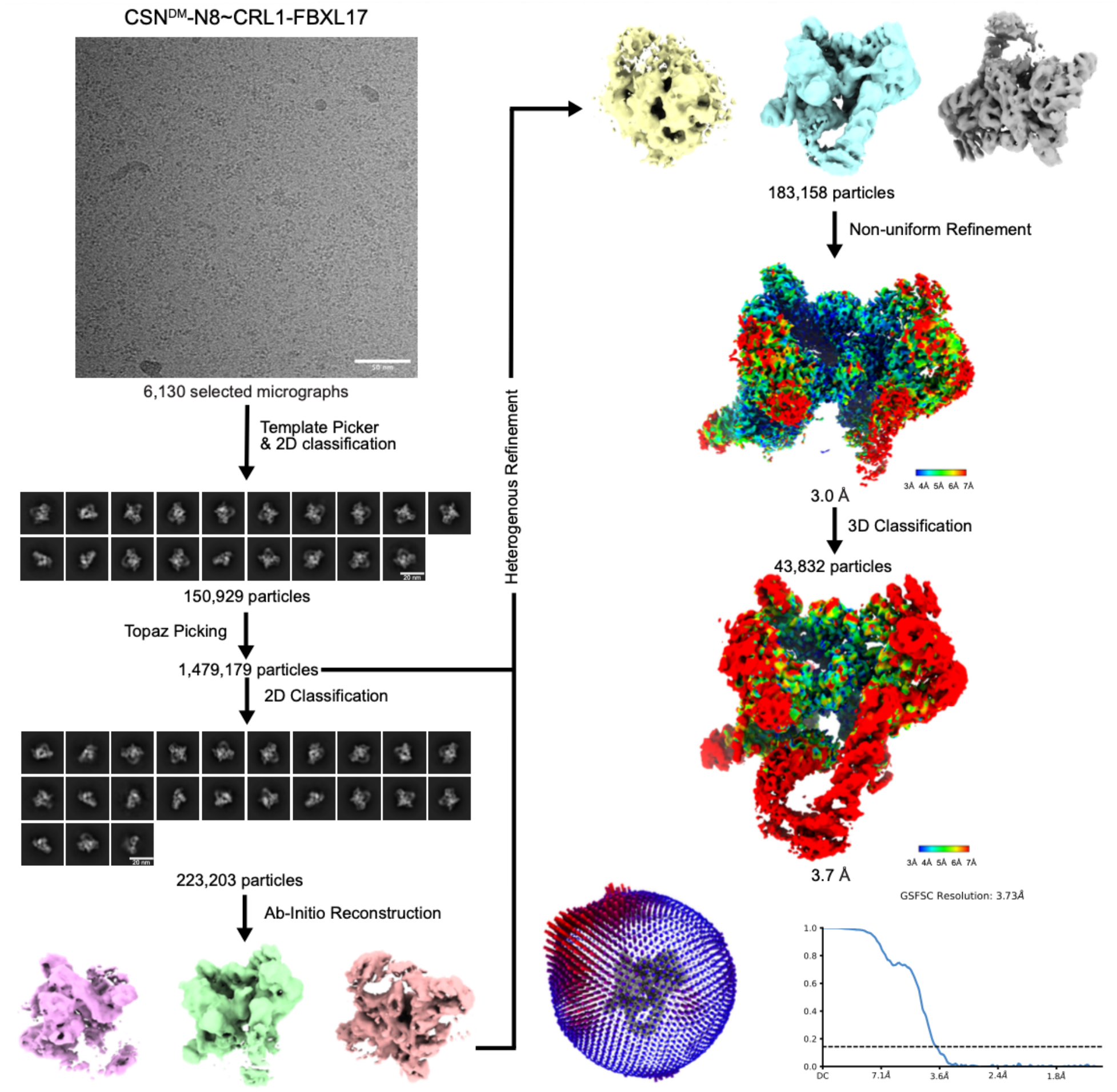
Cryo-EM data processing workflow for CSN^DM^-SCF^FBXL17^

**Supplementary Figure 2.**
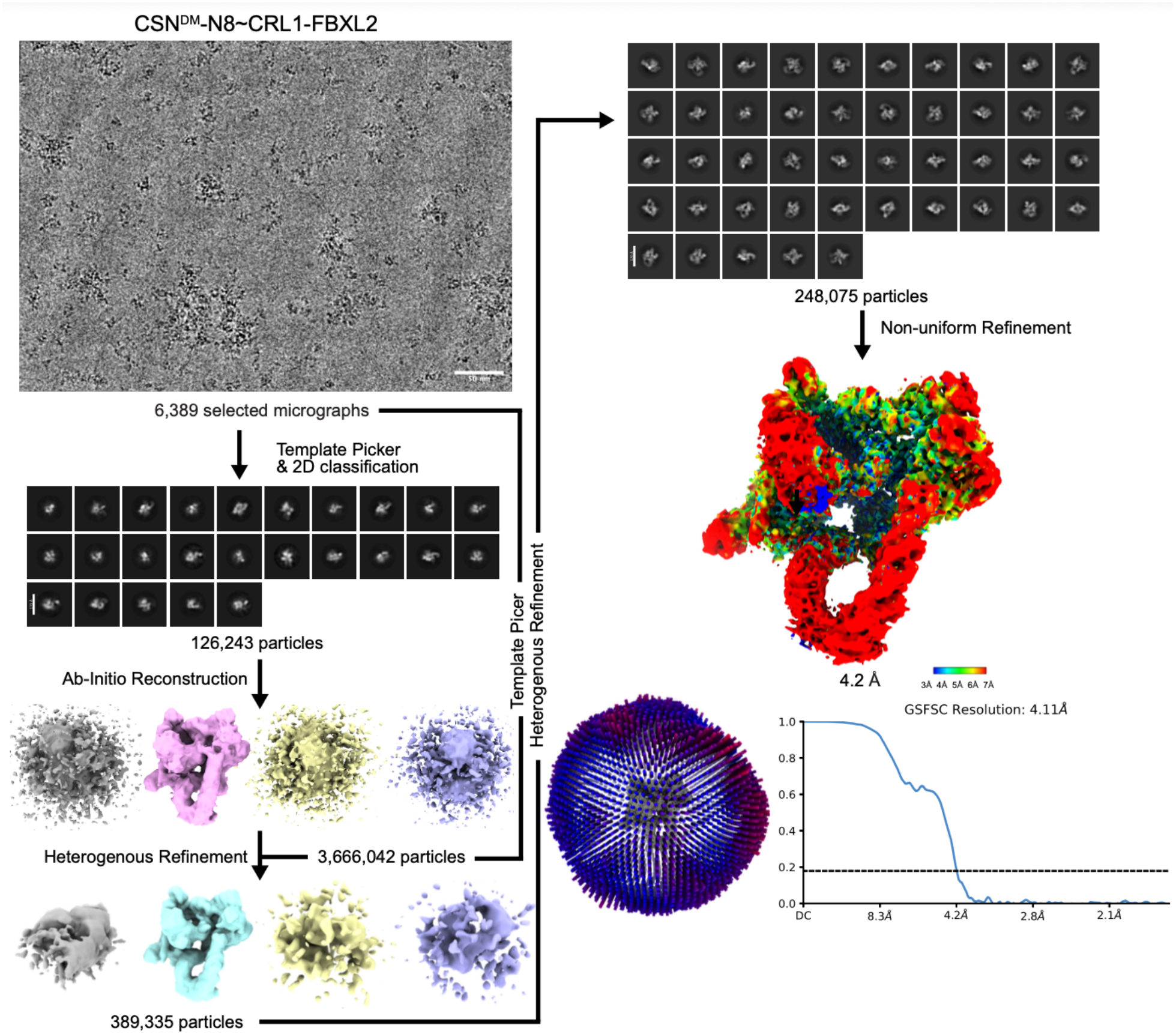
Cryo-EM data processing workflow for CSN^DM^-SCF^FBXL2^

**Supplementary Figure 3.**
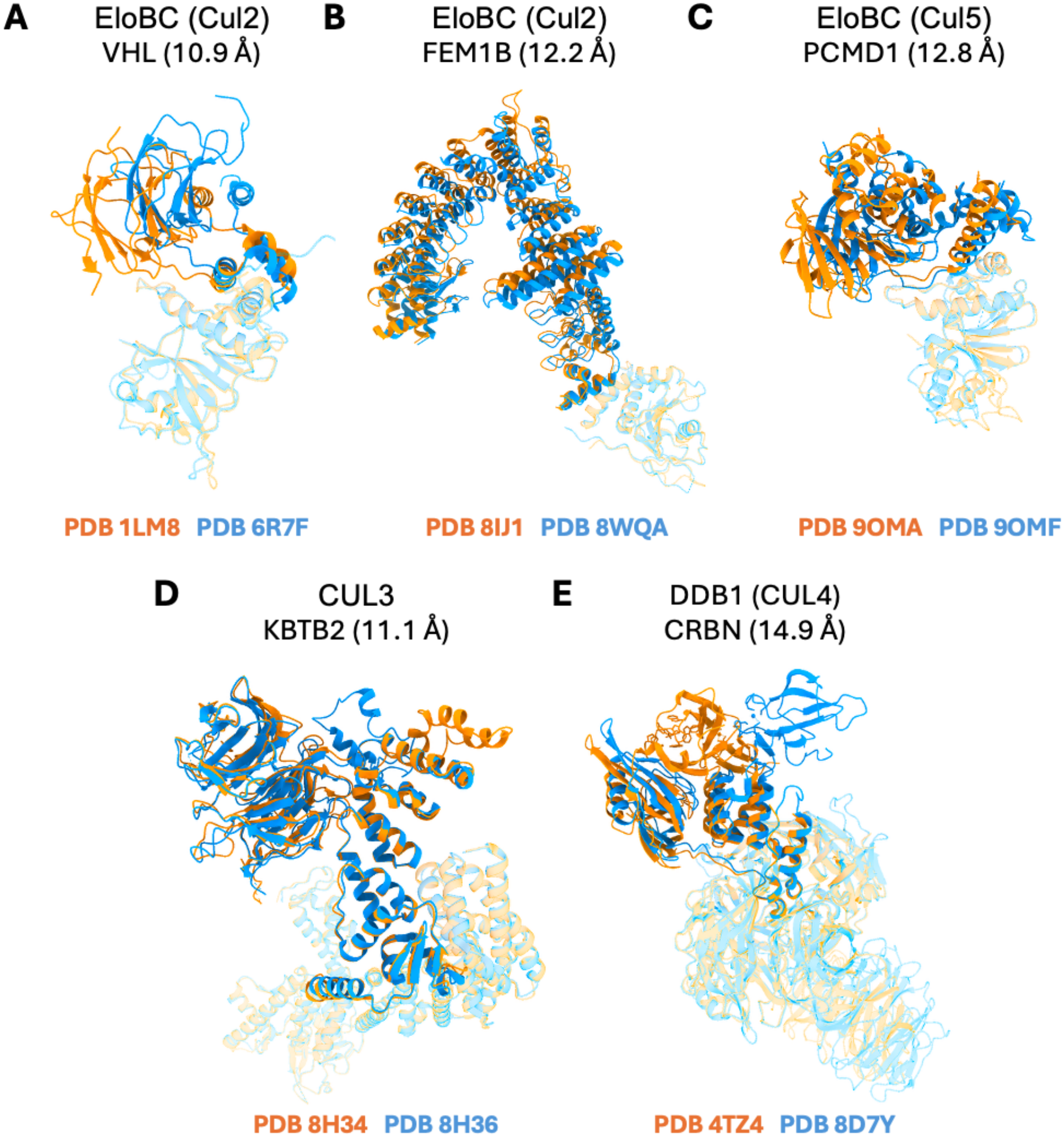
Inter-domain motion across substrate receptors of other CRL families. For five non-CRL1 substrate receptors, two deposited structures were aligned on the adaptor module (partially transparent) to highlight relative motion of the receptor domain (opaque). Each panel reports the receptor-domain Cα RMSD between the two states. **(A)** VHL, EloBC (CUL2) context (PDB 1LM8, orange; 6R7F, blue); 10.9 Å. **(B)** FEM1B, EloBC (CUL2) (8IJ1; 8WQA); 12.2 Å. **(C)** PCMD1, EloBC (CUL5) (9OMA; 9OMF); 12.8 Å. **(D)** KBTB2, CUL3 (8H34; 8H36); 11.1 Å. **(E)** CRBN, DDB1 (CUL4) (4TZ4; 8D7Y); 17.1 Å. As in Figure 1, modules were cropped from their original macromolecular contexts, and alignments use the adaptor subunit positionally analogous to SKP1.

**Supplementary Figure 4.**
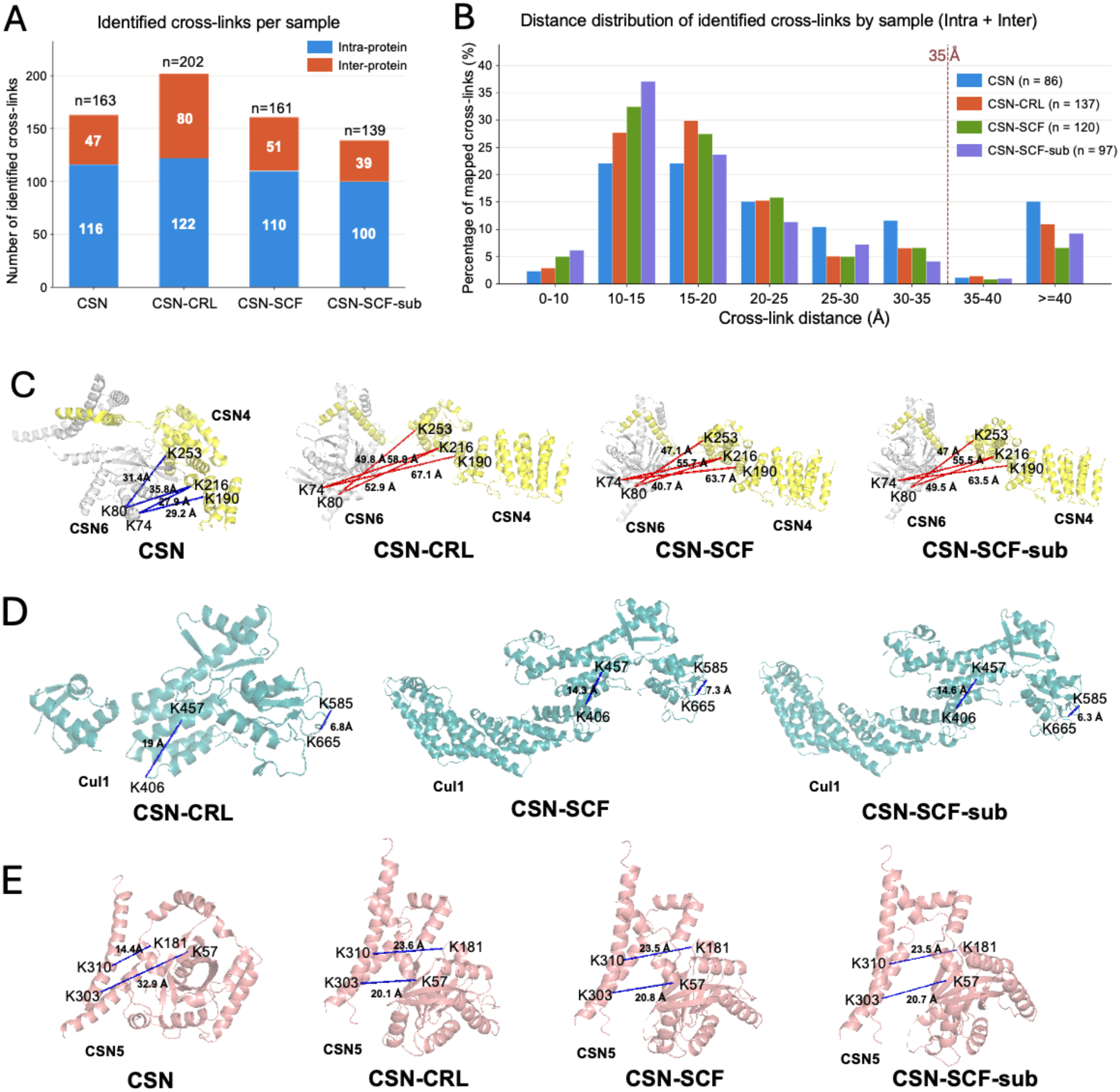
Comparative XL-MS of FBXL2-focused CSN assemblies. **(A)** Number of intra-protein (blue) and inter-protein (orange) DSSO cross-links identified in each sample: CSN, CSN-CRL, CSN-SCF^FBXL2^ and CSN-SCF^FBXL2^-GGTase3. **(B)** Cα-Cα distance distribution of highly reproducible cross-links (identified in ≥6 independent experiments) mapped onto the corresponding structures. **(C)** CSN4-CSN6 cross-links mapped onto each structure. Cross-links that satisfy the constraint are shown in blue and violating distances in red, corresponding to the opening of the CSN4-CSN6 interface upon CRL binding. **(D)** Representative intra-Cul1 cross-links showing minimal distance changes across assembly states. **(E)** Intra-CSN5 cross-links that stay within the distance constraint despite CSN5 conformational changes between assemblies.

**Supplementary Figure 5.**
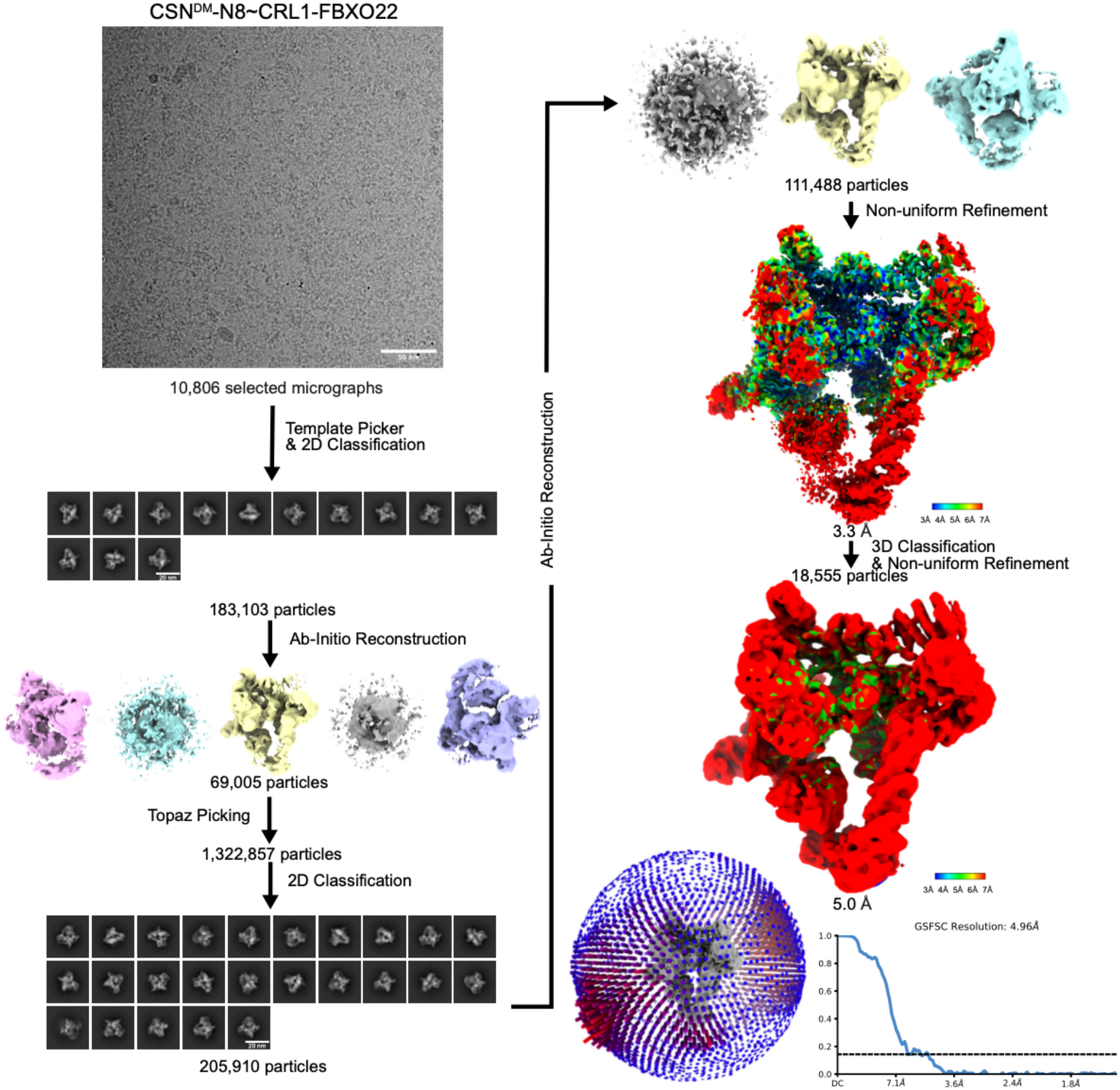
Cryo-EM data processing workflow for CSN^DM^-SCF^FBXO22^

**Supplementary Table 1.** : Cryo-EM image processing and model refinement statistics.

|  | **CSN<sup>DM</sup>-N8~CRL1-FBXL2 (38QO/EMD-79052)** | **CSN<sup>DM</sup>-N8~CRL1-FBXL2-GGTase3 (38SN/EMD-79111)** | **CSN<sup>DM</sup>-N8~CRL1-FBXL17 (38KU EMD-78892)** | **CSN<sup>DM</sup>-N8~CRL1-FBXO22 (38XZ/EMD-79223)** |
|---|---|---|---|---|
| Data collection and processing |  |  |  |  |
| Magnification | 45,000 | 105,000 | 105,000 | 105,000 |
| Voltage (kV) | 200 | 300 | 300 | 300 |
| Electron exposure (e<sup>–</sup>/Å<sup>2</sup>) | 50 | 60 | 60 | 60 |
| Defocus range (μm) | 0.6~2.5 | 0.8~3.0 | 0.8~3.0 | 0.8~3.0 |
| Pixel size (Å) | 0.866 | 0.84 | 0.743 | 0.743 |
| Symmetry imposed | C1 | C1 | C1 | C1 |
| Initial particle images (no.) | 3,666,042 | 986,150 | 1,479,179 | 2,609,705 |
| Final particle images (no.) | 248,076 | 148,964 | 43,832 | 18,555 |
| Map resolution (Å) | 4.2 | 3.9 | 3.7 | 5.0 |
| FSC threshold | 0.143 | 0.143 | 0.143 | 0.143 |
| Map resolution range (Å) | 4.2~10 | 3.9~12 | 3.7~10 | 5~10 |
| Refinement |  |  |  |  |
| Initial model used (PDB code) | 9EFM, 1LDJ, 6O60 | 9EFM, 1LDJ, 6O60 | 9EFM, 1LDJ, 8UBT | 9EFM, 1LDJ, 8UA6 |
| Model resolution (Å) | 4.2 | 3.9 | 3.7 | 5.0 |
| FSC threshold | 0.143 | 0.143 | 0.143 | 0.143 |
| Map sharpening *B* factor (Å<sup>2</sup>) | 135 | 116.6 | 49.6 | 71.6 |
| Model composition |  |  |  |  |
| Non-hydrogen atoms | 32114 | 37247 | 32064 | 32185 |
| Protein residues | 4005 | 4640 | 3993 | 4017 |
| Ligands | ZN:3 | ZN:4 | ZN:3 | ZN:3 |
| *B* factors (Å<sup>2</sup>) |  |  |  |  |
| Protein | 66.69 | 103.83 | 293.22 | 349.40 |
| Ligand | 75.62 | 105.71 | 246.89 | 316.38 |
| R.m.s. deviations |  |  |  |  |
| Bond lengths (Å) | 0.003 (0) | 0.008 (0) | 0.006 (0) | 0.003 (0) |
| Bond angles (°) | 0.704 (0) | 1.015 (0) | 0.803 (0) | 0.769 (0) |
| Validation |  |  |  |  |
| MolProbity score | 1.96 | 1.90 | 2.04 | 2.08 |
| Clashscore | 11.43 | 9.12 | 12.47 | 12.88 |
| Poor rotamers (%) | 0.00 | 1.20 | 0.00 | 0.00 |
| Ramachandran plot |  |  |  |  |
| Favored (%) | 94.40 | 94.91 | 93.26 | 92.70 |
| Allowed (%) | 5.45 | 4.69 | 6.59 | 7.05 |
| Disallowed (%) | 0.15 | 0.39 | 0.15 | 0.25 |

